# PCA-Plus: Enhanced principal component analysis with illustrative applications to batch effects and their quantitation

**DOI:** 10.1101/2024.01.02.573793

**Authors:** Nianxiang Zhang, Tod D. Casasent, Anna K. Casasent, Shwetha V. Kumar, Chris Wakefield, Bradley M. Broom, John N. Weinstein, Rehan Akbani

## Abstract

**Background:** Principal component analysis (PCA), a standard approach to analysis and visualization of large datasets, is commonly used in biomedical research for detecting similarities and differences among groups of samples. We initially used conventional PCA as a tool for critical quality control of batch and trend effects in multi-omic profiling data produced by The Cancer Genome Atlas (TCGA) project of the NCI. We found, however, that conventional PCA visualizations were often hard to interpret when inter-batch differences were moderate in comparison with intra-batch differences; it was also difficult to quantify batch effects objectively. We, therefore, sought enhancements to make the method more informative in those and analogous settings.

**Results:** We have developed algorithms and a toolbox of enhancements to conventional PCA that improve the detection, diagnosis, and quantitation of differences between or among groups, e.g., groups of molecularly profiled biological samples. The enhancements include (i) computed group centroids; (ii) sample-dispersion rays; (iii) differential coloring of centroids, rays, and sample data points; (iii) trend trajectories; and (iv) a novel separation index (DSC) for quantitation of differences among groups.

**Conclusions:** PCA-Plus has been our most useful single tool for analyzing, visualizing, and quantitating batch effects, trend effects, and class differences in molecular profiling data of many types: mRNA expression, microRNA expression, DNA methylation, and DNA copy number. An early version of PCA-Plus has been used as the central graphical visualization in our MBatch package for near-real-time surveillance of data for analysis working groups in more than 70 TCGA, PanCancer Atlas, PanCancer Analysis of Whole Genomes, and Genome Data Analysis Network projects of the NCI. The algorithms and software are generic, hence applicable more generally to other types of multivariate data as well. PCA-Plus is freely available in a down-loadable R package at our MBatch website.

## 1 BACKGROUND

### 1.1 Conventional PCA

Principal component analysis (PCA) [1] is a powerful dimensionality-reduction algorithm for analysis and visualization of multivariate data. It uses linear transformation of data to a new set of variables, principal components (PCs) that are uncorrelated with each other. The PCs are ordered so that the first PC retains as much as possible of the variation present in the original dataset, the second PC retains as much as possible of the remaining variation, and so forth [2]. The PCA algorithm seeks linear combinations of the variables and is sensitive to outliers, but it makes minimal other assumptions about the distribution of the data. Because it is based on linear transformation (unlike UMAP, t-SNE, and other such non-linear dimensionality reduction algorithms), it preserves quantifiable relationships among the locations of the data points. It has been widely used as an exploratory tool for pattern recognition and interpretation of high dimensional data [3] and has been used in a wide variety of contexts, for example, image processing and compression, characterization of molecular dynamics, linguistic information retrieval, and assessment of batch effects [1, 4]. PCA can highlight patterns of important variation and identify groupings of data points in a transparent way. Although the usual PCA visualizations in two or three dimensions capture only a portion of the overall variation in the data, that portion often includes much of the total variation [2]. In biomedical research, PCA is commonly applied to the analysis and visualization of tissue samples that have been profiled for a molecular characteristic such as gene expression, protein expression, DNA copy number, or DNA methylation. It can indicate whether the samples fall into a single, more-or-less homogeneous group based on the profile data or into separate groups. However, conventional PCA is limited in several ways:

1. When the number of samples or number of groupings is large, a conventional PCA plot can fail to distinguish overlapping but distinct groups of sample points.
2. There is no mechanism to portray trends, for example trends in time (or pseudo-time) among groups of data points.
3. The data points are generally color-coded and/or annotated in advance according to pre-existing group or class definitions. It can be hard to distinguish dichotomies or other anomalous distributions in the data if they do not correspond to the pre-existing group definitions. That can make it hard to discover new, influential technical or biological batch variables.
4. Because the principal component projection minimizes squared reconstruction errors, PCA is sensitive to outliers. (The effect of the outliers can be reduced by using robust L1-norm PCA [5], but that is rarely done.)
5. There is no generally applied quantitative index of the global variation among groups of data points.

Here, we present algorithms and a software package, PCA-Plus, that address those limitations of conventional PCA. We propose enhancements that make the plots more informative and introduce a new metric, the dispersion separability criterion (DSC), to quantify global dissimilarities among groups of data. The DSC is accompanied by a permutation test *p*-value for assessing statistical significance.

### 1.2 Applications to The Cancer Genome Atlas (TCGA)

PCA-Plus can be used in any context in which conventional PCA is used. Our motivating examples are found in data from The Cancer Genome Atlas (TCGA) project, a large-scale joint enterprise of the National Cancer Institute and National Human Genome Research Institute in USA [6-10]. TCGA profiled 33 human tumor types at the DNA, RNA, protein, and chromosomal levels over a ten-year period, supporting the dual aims of a better understanding of cancer biology and improved therapy for individuals with cancer. As active members of the TCGA Research Network, we applied PCA-Plus to the TCGA datasets (along with other statistical methods) to assess batch effects and trends in the data. The results were updated every few months and made publicly available through an interactive website. The PCA-Plus visualizations and quantitative analyses successfully detected and diagnosed anomalies in the TCGA data that were then corrected before downstream analyses were performed by the TCGA Disease Working Groups.

In the following section, we describe our methods, including derivation of a computationally efficient formulation for the novel DSC metric and its permutation test *p*-value. In Section 3, we use TCGA data to provide illustrative examples of the features of PCA-Plus, and in Section 4, we state our conclusions. PCA-Plus is freely available in a down-loadable R package at our MBatch website (see Declarations section).

## 2 METHODS

### 2.1 PCA with centroids

As a dimensionality reduction technique, PCA linearly combines optimally weighted observed variables. For clarity, we consider a data matrix with genes (or proteins) in rows and tissue samples in columns. Entries in the matrix may represent gene expression, copy number, methylation, or any other continuous (or ordinal) variable. For use in describing our extensions of PCA, we offer here a brief intuitive explanation of standard PCA and follow up with more technical explanations of PCA and PCA-Plus.

Consider the vector of values for each sample as occupying a point in a space of dimension *n*, where there is one dimension for each gene. On an intuitive level, there are 5 steps in PCA. (i) Center the cloud of points (one for each sample) at the origin (0, 0). (ii) Rotate the orthogonal coordinate system so that one axis (the first principal component axis) explains as much of the variance of the data points as possible. (If the cloud of points were cigar-shaped, that axis would lie along the length of the cigar.) (iii) Rotate the remaining *n*-1 orthogonal axes so that the second principal component axis explains as much as possible of the remaining variance. (iv) Repeat those steps for the remaining *n*-2 axes. (v) Delete the axes that explain the least amount of variance in the data. For visualization, one generally keeps only the first two or three axes.

In mathematical terms: Given a numerical data matrix **Y**_*(N ×D)*_, where *N* is the number of genes and *D* is the number of samples with *N>>D*, the *(i, j)* component of the matrix, *y*_*ij*_, denotes the level of the *i*-th gene in the *j*-th sample, and the row vector ***y***_*i*_ denotes the data vector for the *i*-th gene. The covariance matrix S for vectors ***y***_*i*_ (1 ≤ *i* ≤ *N)* is given by

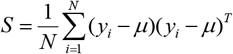

where the vector of the means for the genes is defined as 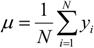, and *T* denotes the transpose of a vector or matrix.

Let *λ*_1_ ≥ *λ*_2_ ≥ · · · ≥ *λ*_*D*_ and ***u***_1_, ***u***_2_, …, ***u***_*D*_ denote the eigenvalues and corresponding eigenvectors of S, respectively. The matrix with eigenvectors corresponding to the largest *L (*1 ≤ *L* ≤ *D)* eigenvalues is **U**=(***u***_1_, ***u***_2_, …, ***u***_*L*_). The first *L* principal components **X** _(*D*×*L)*_ then can be obtained as *X* = *U*^*T*^*Y*, where the column vector ***x***_*l*_ denotes the *l-*th principal component of **Y** and can be calculated as *x* = *Y*^*T*^*u*.

Let **P**_*(K×L)*_ denote the centroids of the L principal components for K groups of samples and let W_*(D×K)*_ denote the design matrix for samples belonging to *K* groups, where

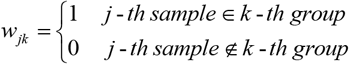

The row vector of centroids for the *k*-th group and *L* principal components ***p***_*k*_ can be obtained by

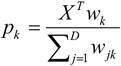

### 2.1 Dispersion separability criterion (DSC): a metric for the global dissimilarity of sets of pre-defined groups or batches

Scatter matrices and metrics derived from them have been described in the literature [11] and used in such contexts as classification [12] and feature selection for unsupervised learning [13]. Here, we introduce a novel variant metric, the *dispersion separability criterion* (DSC), for quantifying the global dissimilarity of sets of pre-defined groups, with application to PCA plots. The DSC can be used, for instance, to assess the magnitude of batch effects or the differences among classes or subtypes of biological samples. Roughly speaking, it assesses the relationship between inter-group variation and intra-group variation in a way that takes into account the relative sizes of the groups. Given an estimated value for the DSC, we use a permutation test to compute *p*-values and confidence limits for that estimate.

The DSC is defined as

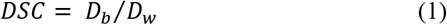

where

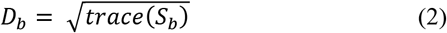

and

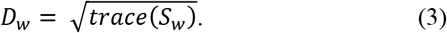

*S*_*b*_ and *S*_*w*_ are the between-group and within-group scatter matrices, respectively (defined in Equations 4–6 below). DSC is the ratio of the average dispersion between group centroids and the average dispersion of samples within groups. A higher DSC value means greater dispersion among groups with respect to that within groups. Defining *D*_*b*_ and *D*_*w*_ as in Equations 2 and 3 enables us to simplify the calculations considerably, making them tractable for large numbers of samples. In addition, the definitions have intuitive appeal in that *D*_*b*_ can be interpreted as the mean distance between group centroids and the global mean, whereas *D*_*w*_ can be approximately viewed as the mean distance between samples within a group and the group centroid.

The within-group scatter matrix, *S*_*w*_, and between-group scatter matrix, *S*_*b*_, are defined as follows [13]:

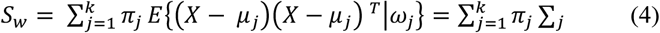

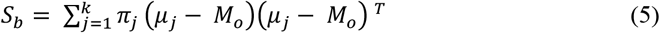

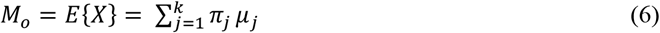

where *π*_*j*_ is the probability that an instance belongs to group *ω*_*j*_, *X* is a *d*-dimensional random feature vector representing the data, *k* is the number of groups, *μ*_*j*_ is the sample mean vector of group *ω*_*j*_, *M*_*o*_ is the total sample mean, ∑_*j*_ is the sample covariance matrix of group *ω*_*j*_, and *E*{.} is the expected value operator.

Equations 4–6 pose a problem for large-volume calculations (such as those we have done over all TCGA datasets): They involve matrix operations that can take considerable compute time and memory. The following derivation leads to a computational form for the DSC that addresses that problem. Substituting Equation 4 into Equation 3, squaring both sides, and simplifying gives us

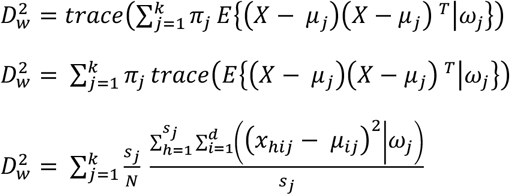

and finally

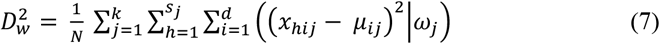

where *N* is the total number of samples in the dataset, *k* is the number of groups, *d* is the number of dimensions in the dataset (e.g., number of genes), *s*_*j*_ is the number of samples in group *ω*_*j*_, *x*_*hij*_ is the value of the *i*^th^ dimension of the *h*^th^ sample belonging to group *ω*_*j*_, and *μ*_*ij*_ is the mean of dimension *i* for all samples in group *ω*_*j*_.

Similarly, substituting Equation 5 into Equation 2, squaring both sides, and simplifying gives us

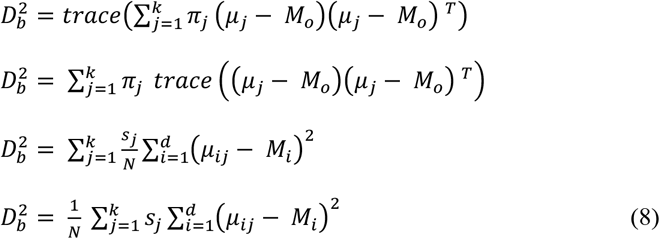

where *M*_*i*_ is the global mean for dimension *i* across all of the samples in the dataset. Note that Equations 7 and 8, unlike Equations 4 and 5, avoid matrix multiplication operations, greatly speeding up the computations and reducing memory requirements. As noted above, that is one of the reasons for defining *D*_*b*_ and *D*_*w*_ as in Equations 2 and 3. DSC can then be computed rapidly by substituting Equations 7 and 8 into Equation 1.

To assess the statistical significance of the DSC metric, we permute the group labels to compute a *p*-value. In each permutation, samples are randomly assigned to groups, keeping the sizes of the groups the same as in the original calculation. The fraction of the time that the permuted DSC value is greater than or equal to the original DSC value provides a nonparametric one-tailed *p*-value with respect to the null hypothesis that the samples are random with respect to group.

### 2.1 Interpretation of the DSC and its *p*-value

The DSC is a continuous positive variable with no obvious threshold value to use in deciding whether the groups are *meaningfully* different. Based on our experience, however, a general rule of thumb is that a DSC value significantly below 0.3 indicates reasonable consistency among groups (i.e., low batch effects); values between 0.3 and 0.6 indicate moderate differences; and values above 0.6 usually indicate strong differences (i.e. strong batch effects). If the DSC is being used to assess batch effects, for instance, and the DSC is greater than 0.6 the data may need to be corrected for batch effects before they can be used for analysis or prediction.

As in other parameter-estimation contexts, both the DSC estimate and its *p*-value must be considered. Outlier samples can lead to a misleadingly large value of DSC but non-significant *p*-value, particularly if the numbers of samples per group are small. Conversely, the calculated value of the DSC may be small but the *p*-value significant if the numbers of samples per group are large.

## 3 RESULTS

This section describes enhancements over standard PCA (in addition to the DSC metric and *p*-value) that we have incorporated into PCA-Plus. Most of the examples presented here relate to the assessment of batch effects in TCGA datasets. However, PCA-Plus is much more generic; it can be applied to any context in which the samples are preassigned to particular groups. One such context, comparison of different tumor subtypes, is described in Section 3.5. Whereas we generally hope to observe minimal differences among groups in analysis of technical batch-effects, we are usually pleased to find large differences when comparing biological classes or subtypes of samples. PCA-Plus can also be used when the roles of genes and samples are reversed — for example, when assessing the expression levels of groups of genes that belong to different pathways or different functional categories in the Gene Ontology database.

### 3.1 Batch-effects analysis of TCGA data

A brief description of batch effects in data from TCGA and analogous molecular profiling projects will be useful to the interpretation of the examples that follow. Technical (i.e., non-biological) batch effects could arise at any number of levels in TCGA data processing. The project acquired tumor and normal tissue samples from dozens of institutions designated as tissue source sites. Those sites may have differed in their procedures or in biologically meaningful variables such as the patient populations they served and therefore the characteristics of the samples they accrued. The samples were then processed in one of two Biospecimen Core Resource (BCR) centers, which sometimes used different protocols. The samples were aliquoted by the BCR and sent in batches to the Genome Sequencing Centers and Genome Characterization Centers, where they were processed in batches to generate molecular data at the DNA, RNA, protein, and chromosomal levels. Technical batch effects (or trends over time) could be introduced at any of those levels. PCA-Plus can detect, diagnose, and quantify such effects. Equally important, it can help with domain-aware detective work to identify new definitions of *batch* that had not been considered before the analysis (as in Figures 3 and 4 below).

### 3.2 PCA with centroids

Figure 1(a) shows a conventional two-dimensional PCA plot (first and second principal components) for the TCGA colon adenocarcinoma gene expression (Agilent) dataset. Figure 1(b) shows the same plot but with batch centroids and rays extending from the centroids to within-batch data points. It is difficult to see from Figure 1(a) whether any batch stands out from the others, but Figure 1(b) makes it obvious that batch 29 is distinctive (centroid toward the bottom right of the graph). Centroids visually enhance our ability to spot common types of differences among batches.

**Fig. 1.**
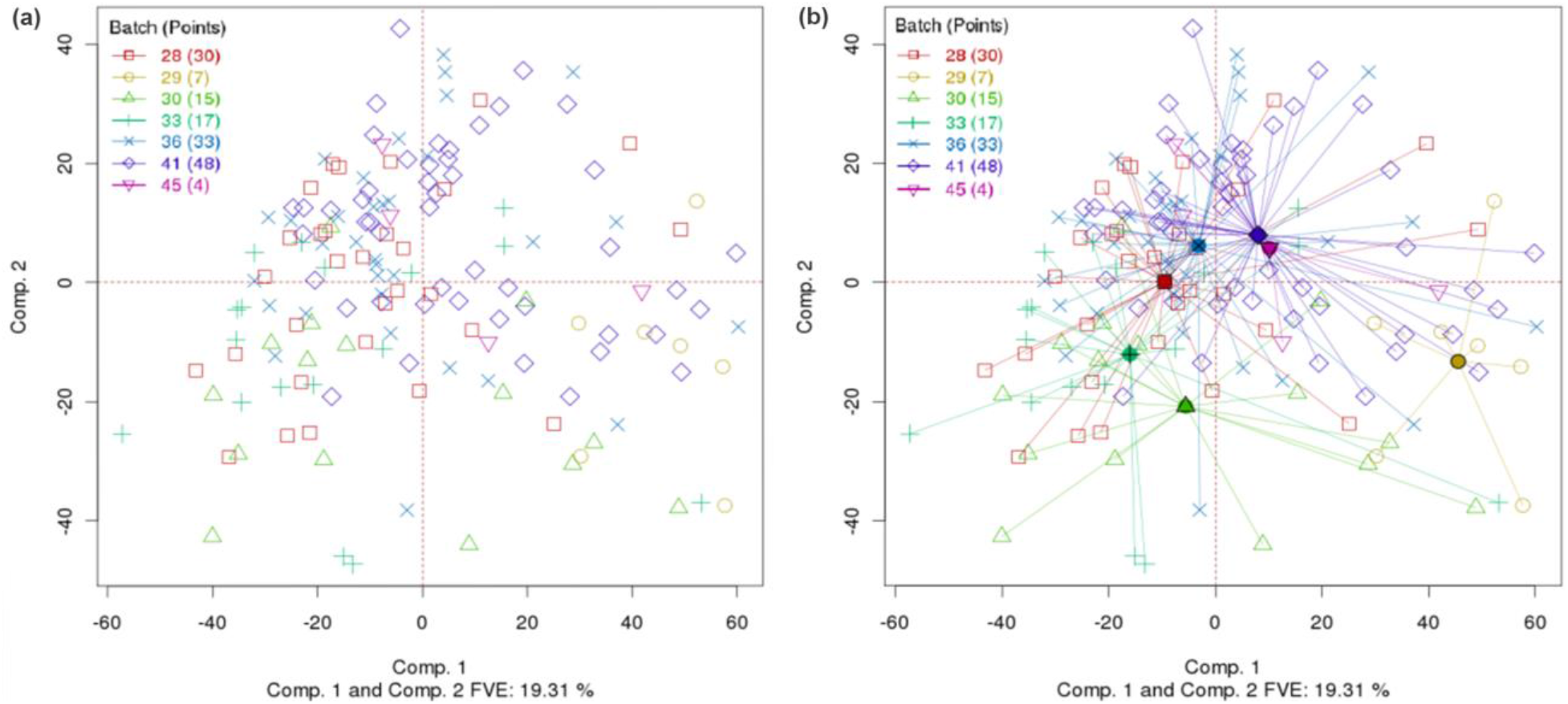
(a) Conventional PCA plot for TCGA colon cancer gene expression (Agilent) data, with points colored by batch ID of the samples. (b) Same PCA plot, but with batches connected by centroids. (DSC for PCA=0.6, p-value < 0.0005; Overall DSC=0.357, p-value < 0.0005).

### 3.3 Trend lines

When the batch variable is ordinal, for example on the basis of time, trajectories (trends) can be indicated by connecting the batch centroids with arrows. If the arrows tend to point in a consistent direction, there may be a trend in the data. Figure 2(a) shows the same data set as in Figure 1, but with batches connected by trajectory lines (black arrows) in order of ascending batch number (and therefore time of processing). No major trend is evident. On the other hand, Figure 2(b) shows the TCGA rectal adenocarcinoma gene expression (Agilent) data set, with ship date (the date the sample was shipped to the BCR) as the batch variable. A general trend can be seen going from the top right (earliest batch) of the figure towards the bottom left (latest batch), indicating the possibility of a slight drift in the data over time. Fortunately, however, TCGA datasets generally do not show strong trends based on time.

**Fig. 2.**
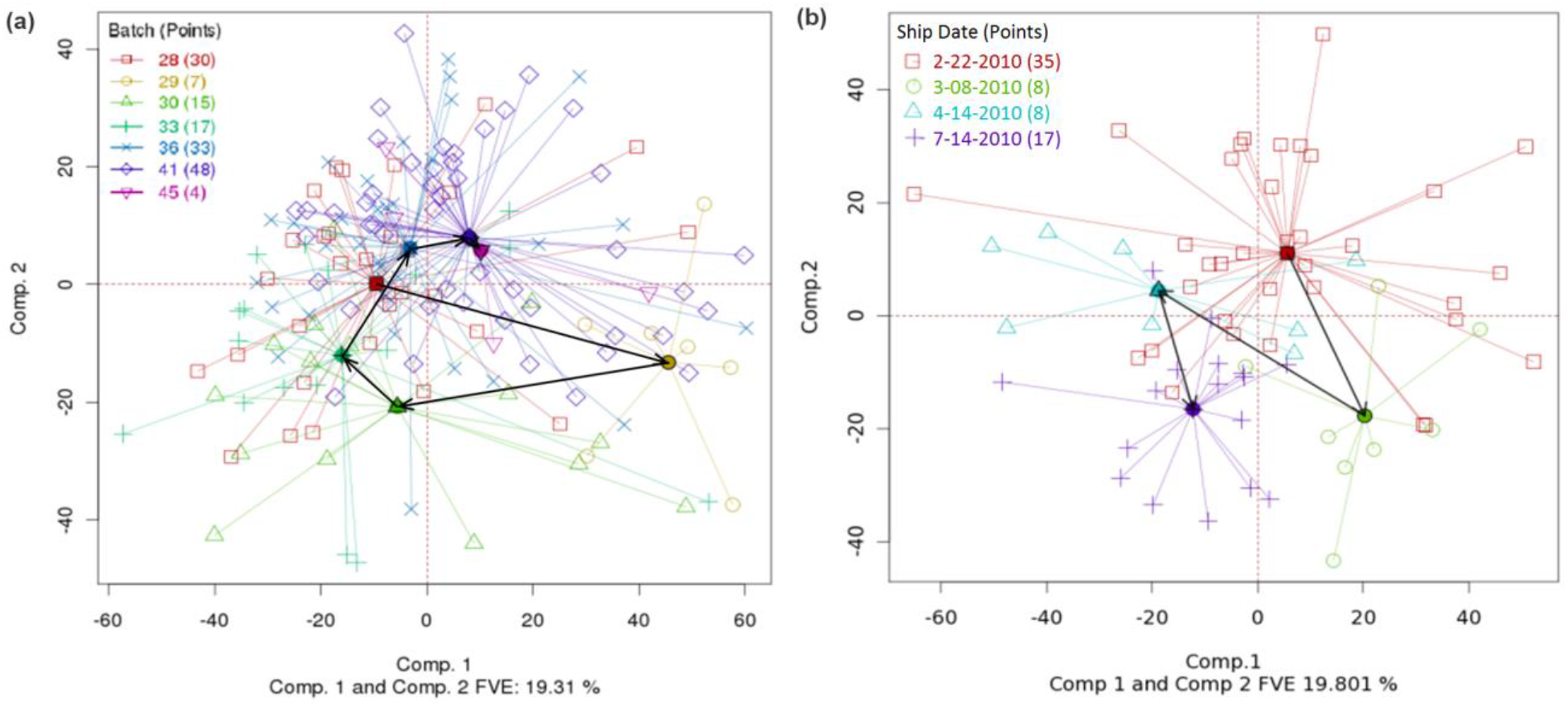
(a) Same data and PCA plot as in Fig. 1, but with centroids connected by trend lines (black arrows). (DSC for PCA=0.6, p-value < 0.0005. Overall DSC=0.357, p-value < 0.0005). (b) TCGA rectal adenocarcinoma (Agilent) gene expression dataset with batch centroids and trend lines. (DSC for PCA=0.706, p-value < 0.0005. Overall DSC=0.344, p-value < 0.0005)

### 3.4 Observation of dichotomies

PCA-Plus makes it easy to spot dichotomies in the data, even if the dichotomies are not based on pre-existing ways of grouping the samples. Figure 3(a) represents TCGA kidney clear cell carcinoma DNA methylation data, with centroids and color coding based on batch ID. There is a clear dichotomy that cannot be attributed to batch ID. Further detective work revealed that the dichotomy was due to differences in the methylation profiles of male and female patients, as seen in Figure 3(b). Removing methylation data for sex chromosomes alleviated the dichotomy. PCA-Plus is generally better than PCA itself for highlighting batch effects that are not based on the pre-existing definitions of *batch*. Interestingly, one patient that was originally labeled female in the clinical data but was observed to group with the male cluster in a previous iteration of the PCA-Plus plot (not shown) was later identified to be male. An error had been made in recording the patient’s sex in the clinical data, which was corrected later. That example illustrates how PCA-Plus can be used to potentially catch such errors.

**Fig. 3.**
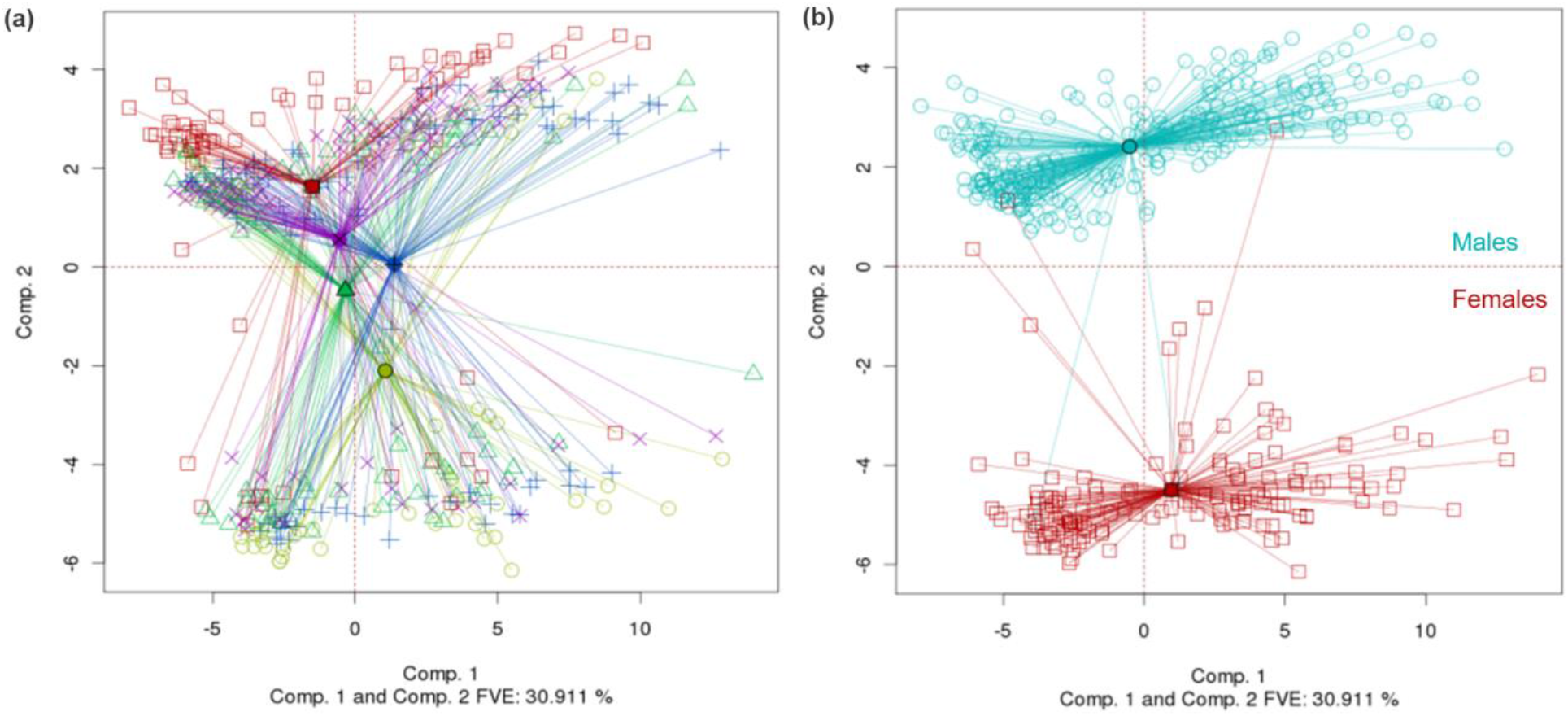
(a) TCGA kidney clear cell carcinoma DNA methylation dataset showing a dichotomy in the data that does not correspond with batch ID. (DSC for PCA=0.267, p-value < 0.0005. Overall DSC=0.334, p-value < 0.0005). (b) Same dataset except the data points have been grouped by patient sex instead of batch ID. The dichotomy corresponds with male vs. female patients. (DSC for PCA=0.706, p-value < 0.0005. Overall DSC=0.341, p-value < 0.0005)

### 3.5 Plotting dual batch variables

The dichotomy described in Section 3.4 can be visualized in another way by comparing Figure 4 with Figure 3(a). In Figure 4, the samples are grouped by batch ID, and the centroids and rays are colored accordingly, as in Figure 3(a). However, the shapes and colors of the points are based on gender as in Figure 3(b). Male and female patients form distinct groups. The PCA-Plus software package enables the user to visualize other combinations of dual variables in an analogous way – for example, patients with good and poor prognosis vs. tumor stage, or batch ID vs. tissue source site, etc.

**Fig. 4.**
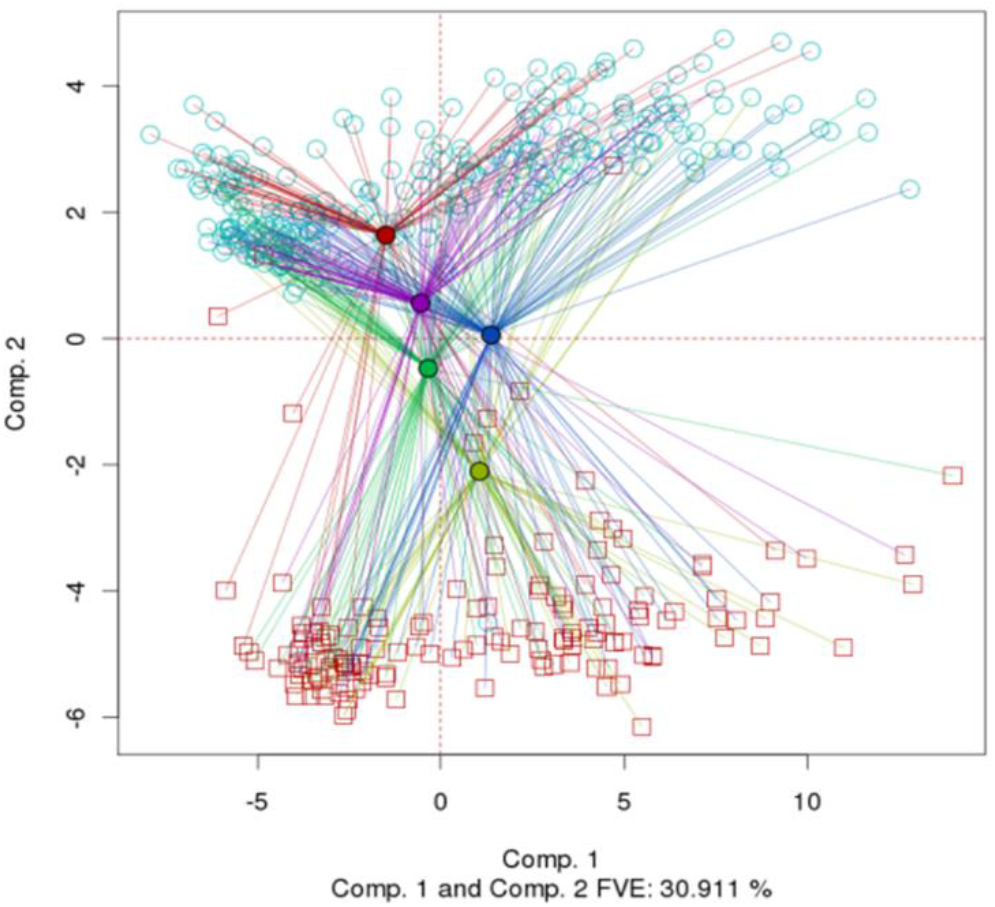
Same dataset as in Fig. 3, but with dual batch enhancement. The rays are colored by batch ID, whereas the points are colored by patient sex. (DSC for PCA=0.267, p-value < 0.0005. Overall DSC=0.334, p-value < 0.0005)

### 3.6 Visualization of subtypes

PCA-Plus can be used to visualize different tumor types or subtypes. Figure 5(a) shows the TCGA glioblastoma multiforme (GBM) gene expression dataset [6], with centroids, rays, and color coding based on subtypes defined by TCGA. The subtypes are largely distinct with a high overall DSC value of 0.514, suggesting large differences in biology between the subtypes. By contrast, Figure 5(b) shows analogous TCGA data for ovarian cancer gene expression-based subtypes, as defined by TCGA [7]. The figure shows that the ovarian subtypes are visually not as distinct as GBM subtypes and there are large areas of overlap between the subtype clusters. The overall DSC value is 0.313, suggesting only moderate differences in biology between the subtypes.

**Fig. 5.**
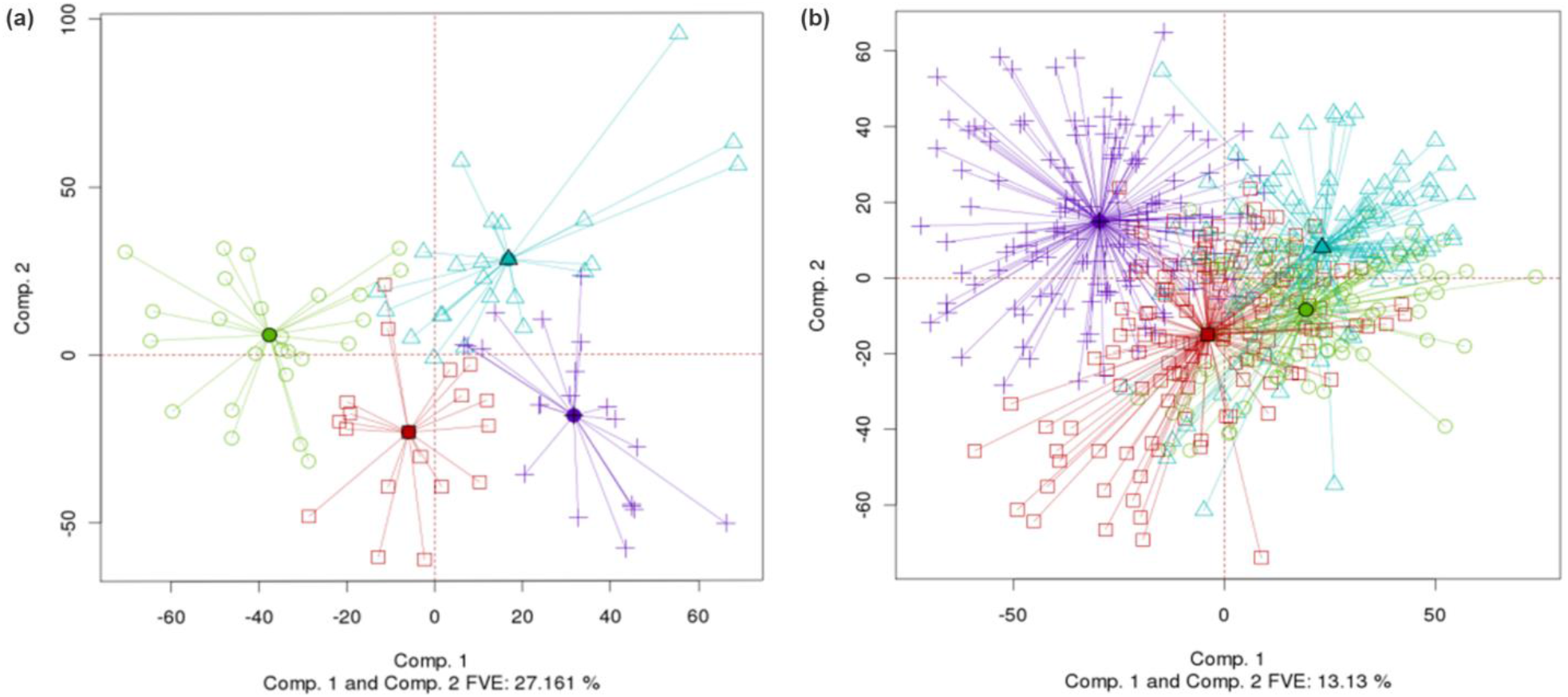
(a) Different subtypes for TCGA glioblastoma multiforme gene expression (Agilent) dataset. (DSC for PCA=1.233, p-value < 0.0005. Overall DSC=0.514, p-value < 0.0005) (b) Different subtypes for TCGA ovarian gene expression dataset. (DSC for PCA=0.905, p-value < 0.0005. Overall DSC=0.313, p-value < 0.0005)

### 3.7 Visualization and assessment of batch effects before and after computational correction

Any of the published correction algorithms can be used to mitigate batch effects identified through PCA-Plus analysis. Figure 6 shows the effect of correcting for batch effects using Empirical Bayes as in ComBat [14]. Figure 6(a) shows uncorrected TCGA gene expression data from rectal adenocarcinoma samples grouped by batch ID. The overall DSC value is 0.356, indicating moderate batch effects in the data. Figure 6(b) shows the same data after correcting for batch effects. The batches appear much more homogeneous, and the DSC value is much lower (0.071), indicating negligible batch effects. Thus, PCA-Plus is useful for quantifying the effectiveness of batch effect correction algorithms.

**Fig. 6.**
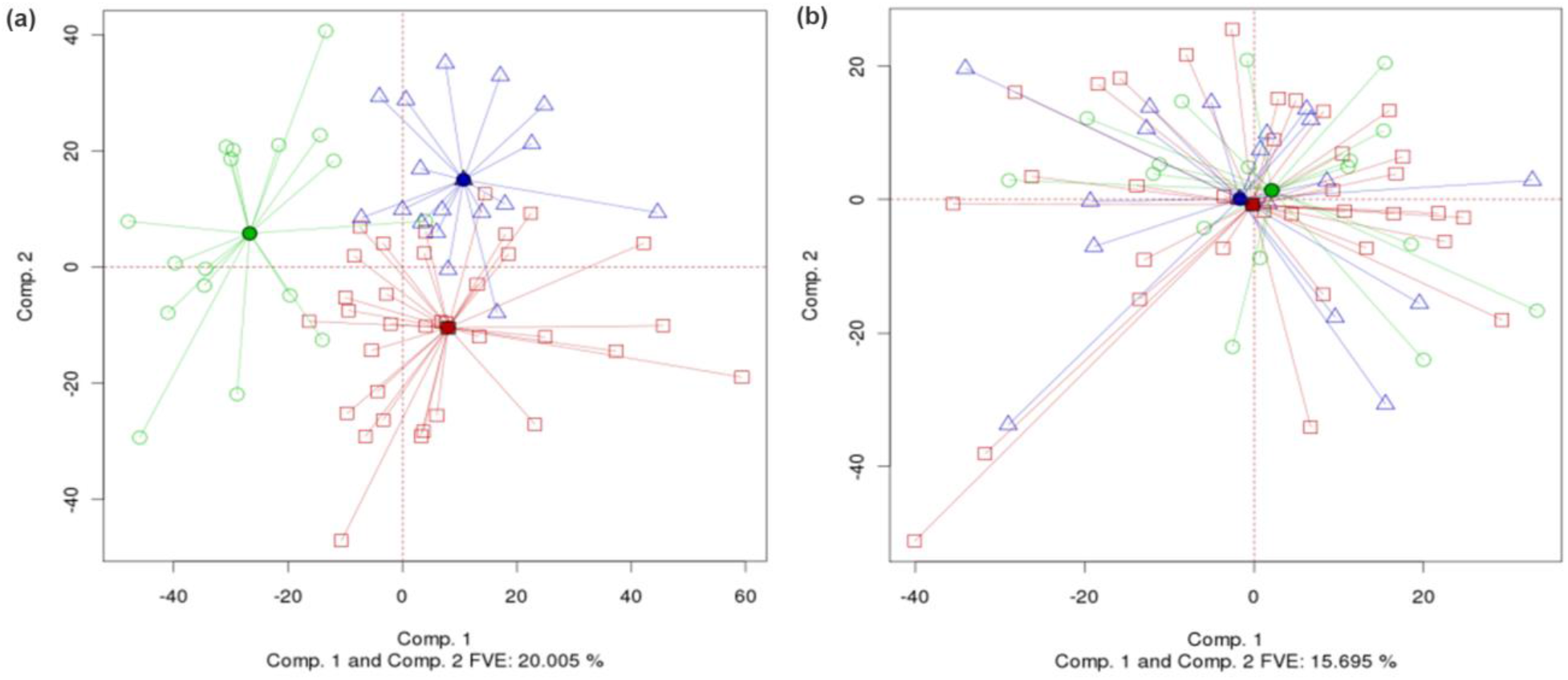
(a) TCGA rectal adenocarcinoma gene expression dataset before correcting for any batch effects. (DSC for PCA=0.896, p-value < 0.0005. Overall DSC=0.356, p-value < 0.0005) (b) Same dataset after using the empirical Bayes algorithm to correct for batch effects. (DSC for PCA=0.069, p-value=0.958. Overall DSC=0.071, p-value=1)

## 4 DISCUSSION

PCA-Plus has been used in all 33 TCGA disease working groups and more than 30 PanCanAtlas groups to detect, diagnose and quantify batch effects in “omics” data. Whenever batch effects were discovered, they were either corrected at the source, e.g. by re-running raw data through an updated software pipeline, or by algorithmic correction down the line. However, algorithmic batch effects correction can often over-correct the data, removing actual biological differences along with technical artifacts. There is a trade-off. By using a biological variable (instead of batch ID) to define batches and computing its DSC value, PCA-Plus can be used to determine whether known biological subtypes tend to maintain their distinctions rather than being adjusted away. A high DSC value for the biological variable (e.g. greater than 0.6), coupled with a low DSC value for the batch variable (e.g. less than 0.3) is desirable. Parameters of the correction algorithm used can be tweaked, the data re-corrected, and patterns in the data then re-analyzed by PCA-Plus to optimize the tradeoff.

## 5 CONCLUSIONS

Standard PCA is a favorite algorithm for comparing intra- and inter-group differences in multivariate data, and the enhanced PCA (PCA-Plus) described here adds a number of useful features: (i) group centroids; (ii) trend arrows (when pertinent); (iii) separate coloring of centroids, rays, and data points; and (iv) quantitation in terms of the new DSC metric with corresponding permutation test *p*-values. Those features greatly enhance one’s ability to visualize and quantitate even modest differences among the groups represented or to conduct detective work on dichotomies that show up but do not correspond to pre-defined batches in the dataset. The algorithms are general, but we have illustrated them here through their application to the detection and quantitation of batch effects, trend effects, dichotomies, and tumor subtype differences in TCGA molecular profiling data. The MBatch software package, which features PCA-Plus, includes modules that generate figures and metrics, as well as hierarchical clustering and other plots that can be used in tandem with PCA-Plus.

## 6 LIST OF ABBREVIATIONS

BCR: Biospecimen Core Resource
DSC: Dispersion Separability Criterion
FVE: Fraction of Variance Explained
GBM: Glioblastoma Multiforme
GDC: Genomic Data Commons
PC: Principal Components
PCA: Principal Component Analysis
TCGA: The Cancer Genome Atlas

## 7 DECLARATIONS

### Ethics approval and consent to participate

All the data used were obtained from publicly available sources. TCGA data were obtained from NCI’s public website and no additional data were generated by the authors.

### Consent for publication

Not applicable

### Availability of data and materials

The datasets analyzed during the current study are available from the Genomic Data Commons (GDC) under TCGA project at https://portal.gdc.cancer.gov/. The software used is freely available from https://bioinformatics.mdanderson.org/public-software/mbatch/. A compendium of PCA-Plus plots we have produced is available at http://bioinformatics.mdanderson.org/tcgabatcheffects.

### Authors’ contributions

NZ, RA and JNW developed the visualizations. RA developed the DSC metric. TDC, CW and BMB wrote the MBatch code. AKC, NZ and RA performed the analyses. NZ, RA, AKC, SVK, BMB and JNW interpreted the results. RA, NZ, AKC, SVK, and JNW wrote and edited the manuscript. All authors reviewed the manuscript.

## Acknowledgements

Not applicable.

## Authors’ information

Authors RA and JNW were principal investigators on a TCGA genome data analysis center (GDAC) grant (NCI U24 CA143883) that focused on batch effects analyses of TCGA and PanCanAtlas data. Early versions of PCA-Plus were used as part of those analyses and they were instrumental in publishing TCGA’s 60+ papers. RA, BMB, and JNW are currently PIs on another grant (NCI U24 CA264006) for batch effects analysis of data from NCI’s Center for Cancer Genomics (CCG), where PCA-Plus is also extensively used.

**Project name:** PCA-Plus

**Project home page:** https://bioinformatics.mdanderson.org/public-software/mbatch/

**Operating system(s):** Platform independent

**Programming language:** R (open-source)

**License:** GNU GPL

**Non-Academic Use:** No restrictions

